# An approach towards identifying the site of interaction between the *Streptomyces* phage ØC31 protein Gp3 and the coiled-coil domain of its serine integrase

**DOI:** 10.1101/2022.09.24.509315

**Authors:** Monalissa Halablab, Sean Colloms, Steven Kane

## Abstract

*Streptomyces* phage ØC31 encodes a serine integrase which recombines the host (attB) and phage (*attP*) attachment sites to form new *attL* and *attR* sites. For *attL* and *attR* to be recombined by the integrase, the presence of its cognate recombination directionality factor (RDF) is required. It is hypothesized that the RDF binds the coiled-coil domain of the integrase to accomplish this switch in recombination directionality. Yet, nothing is known about the site of interaction between the RDF and integrase. Here, we tried to identify the region on the coiled-coil domain of ØC31 integrase to which its cognate RDF (gp3) might bind. Mutant integrases fused to their cognate RDF (gp3) were created and selected for inactivity in *attL* x *attR* recombination. It was not possible to characterize the mutants and identify the Int-RDF binding region, due to unforeseen errors that occurred during the construction of the mutant library, but we demonstrated that simple experimental approaches could be used to identify this region. Despite this, an integrase mutant (P398L D595N) fused to gp3 was characterized. This mutant was catalyzing a more unidirectional *attL* x *attR* recombination reaction with reduced *attP* x *attB* recombination compared to the wild-type integrase. This mutant was selected since it could be used within the field of molecular biology to construct inversion switches which are a key element by which cells can be computerized.

By the time this work was uploaded on bioRxiv, there is now a publication that investigated the Int-RDF interaction interface and several residues at the base of the coiled-coil that affected both the interaction and recombinase activity were identified (Paul C M Fogg, Ellen Younger, Booshini D Fernando, Thanafez Khaleel, W Marshall Stark, Margaret C M Smith, Recombination directionality factor gp3 binds PhiC31 integrase via the zinc domain, potentially affecting the trajectory of the coiled-coil motif, Nucleic Acids Research, Volume 46, Issue 3, 16 February 2018, Pages 1308-1320, https://doi.org/10.1093/nar/gkx1233).

## INTRODUCTION

The integrase (Int) from *Streptomyces* temperate phage ØC31, which has been originally described by Lomovskaya and colleagues (1972), belongs to the serine recombinase superfamily. This enzyme catalyze site-specific integration of the phage genome into bacterial host chromosome during the phage’s lysogenic lifestyle, and excision of its genome out of the bacterial host’s DNA during its lytic cycle (Rowley et al., 2008; Rutherford and Van Duyne, 2014). During integration, Int recombines the host (*attB*) and phage (*attP*) attachment sites, to form new sites *attL* and *attR*, and recombines *attL* and *attR* to regenerate *attB* and *attP* during excision. However, recombination of *attL* and *attR* sites only occurs if a phage encoded protein; recombination directionality factor (RDF) is present (Khaleel et al., 2011; Rowley et al., 2008; Rutherford and Van Duyne, 2014). An ØC31 RDF, gp3, was identified by Khaleel et al. (2011), whereby this excision-stimulating protein, activated *attL x attR* recombination and inhibited *attP x attB* recombination. Therefore, in the absence of RDF, the *attP* x *attB* recombination reaction catalyzed by the Int is unidirectional and site-specific (Fogg et al., 2014; Rutherford and Van Duyne, 2014). Due to the site-specificity and unidirectional nature of the reactions it catalyzes, ØC31 Int has been widely used as a tool for genome engineering (Fogg et al., 2014; Groth and Calos, 2004). It has been used for assembly of metabolic pathways (Colloms et al., 2014), stable genetic engineering of actinomycetes through constructed plasmids that utilize the ØC31 integration system such as pSET152 (Bierman et al., 1992; Baltz, 2012), gene therapy (Chavez and Calos, 2004), logic gates to control transcription rates (Bonnet et al., 2013) and many other various applications (Groth and Calos, 2004; Fogg et al., 2014).

Serine integrase consists of around 130 residue N-terminal catalytic domain conserved across the serine recombinases and a 300-500 residue C-terminal catalytic domain that contains the enzyme’s DNA-binding functionality, connected to each other by a helix (Smith, 2015; Van Duyne and Rutherford, 2013; Rutherford and Van Duyne, 2014). The C-terminal domain is comprised of a recombinase domain and a zinc-ribbon domain that bind DNA with high-affinity (McEwan, Rowley and Smith, 2009) and an extended coiled-coil motif embedded in the zinc domain that controls directionality of recombination (Rowley et al., 2008; Rutherford et al., 2013; Rutherford and Van Duyne, 2014) (Figure 1).

**Figure 1.**
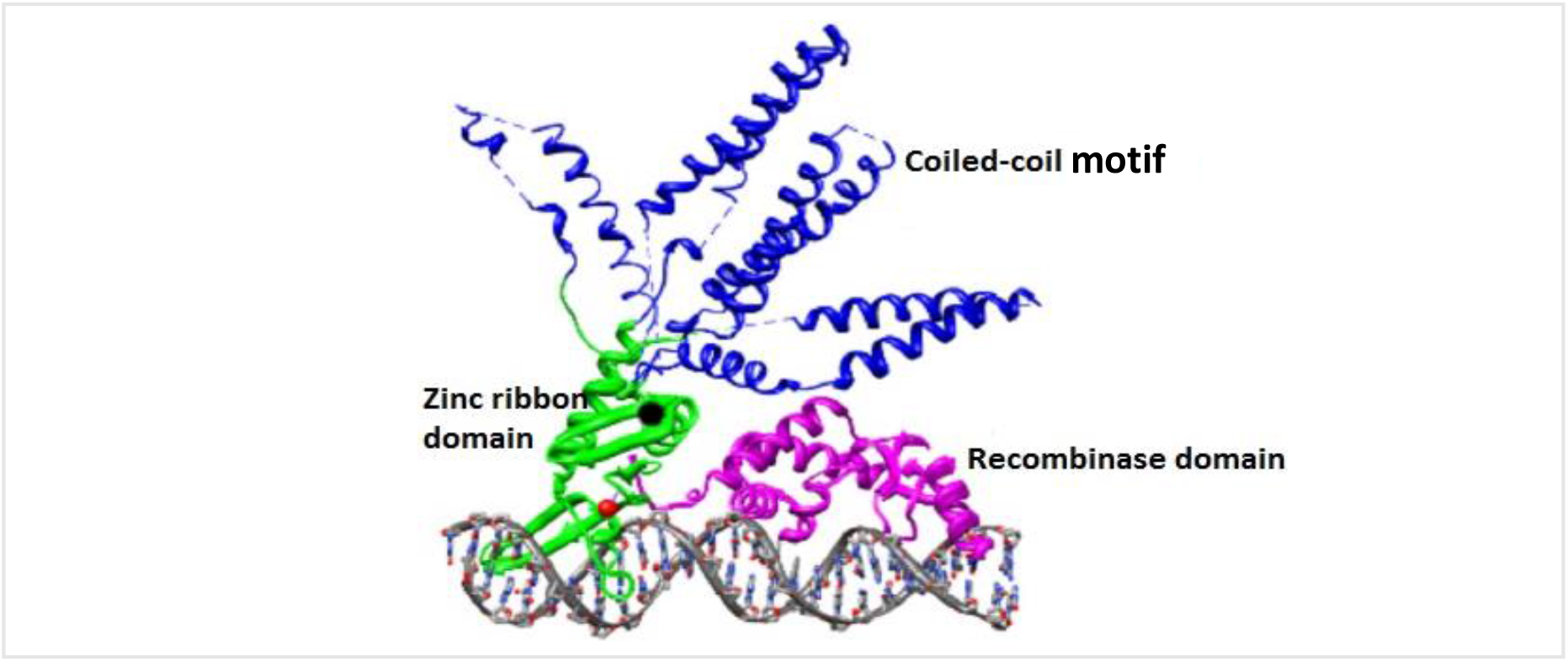
Structure of the C-terminal catalytic domain of Int bound to an attP half-site. The recombinase domain (purple) and zinc ribbon domain (green) are bound to the DNA. The coiled-coil motif (blue) embedded in the zinc domain does not bind DNA (Figure from Rutherford and Van Duyne, 2014).

The current knowledge on the catalytic mechanism of serine integrases has come from the studies done on the “small serine recombinases” such as Tn3 resolvase and Hin invertase (Stark, 2014; Smith, 2015). A model that explains the catalysis mechanism of serine integrases and possible reasons for the unidirectional nature of *attP* x *attB* recombination was proposed by Van Duyne and Rutherford (2013). During integration, the serine integrase binds as a dimer to specific sequences (attachment sites) in the phage (*att*P) and host (*att*B) DNA, mediates association of the sites to form a tetrameric complex and catalyzes recombination of the attachment sites, *attB* and *attP*, to generate new hybrid products *attL* and *attR* (Rutherford and Vun Duyne, 2014). The generated *attL* and *attR* sites do not recombine due to the inhibitory interactions present on Int-*attL* and Int-*attR* by the coiled-coil motifs of the Int (Figure 2) (Rutherford and Vun Duyne, 2014; Thorpe, Wilson and Smith, 2000), resulting in the unidirectional nature of the *attP x attB* recombination reaction. However, only when the RDF is present, *attL and attR* are recombined by Int, with some level of *attP* and *attB* recombination still occurring (Olorunniji et al., 2017). It was also shown, that the RDF (gp3) can be fused to the C-terminus of serine integrase to create single proteins that catalyze *attL* x *attR* recombination with reduced *attP* x *attB* recombination compared to Int and RDF proteins present separately (Olorunniji et al.,2017). However, how RDFs accomplish this switch in directionality is not yet understood. It is hypothesized that the RDF binds to the CC motifs of the serine integrase (Figure 2) in a way that disrupts the inhibitory interactions present on Int-*attL* and Int-*attR*, allowing the formation of stable synaptic complexes and recombination of *attL* and *attR* (Rutherford and Van Duyne, 2014).

**Figure 2.**
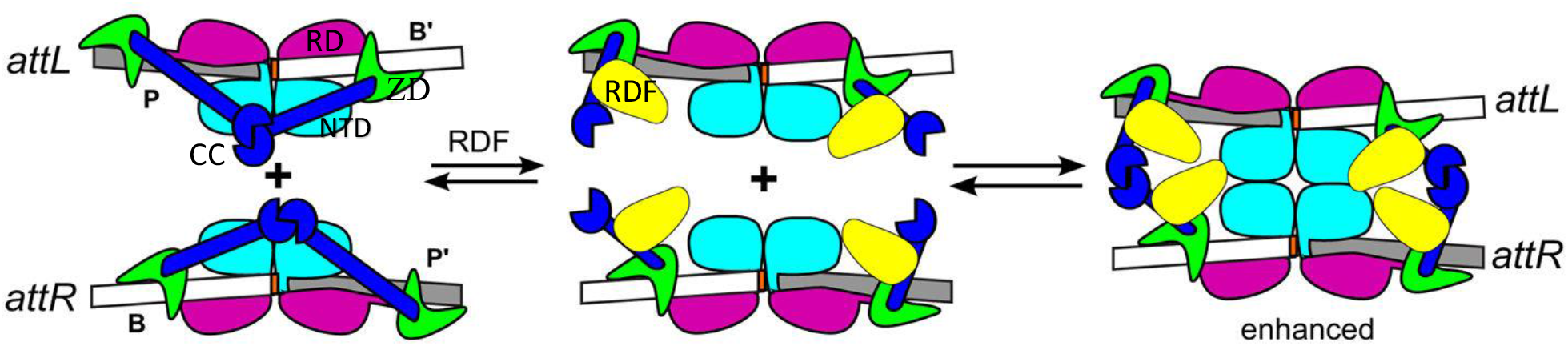
Proposed model for the RDF-Integrase excision mechanism. The RDF (yellow) binds to the CC motifs (blue) of the integrase and releases the inhibition on *attL x attR* recombination. NTD: N-terminal catalytic domain (cyan); RD: recombinase domain (purple); ZD: zinc ribbon domain (green); CC: coiled-coil motif (blue). (Figure from Rutherford and Van Duyne, 2014)

These current hypotheses are based on mutational analysis and research that has been conducted on the CC region of ØC31 integrase since it plays a key role in integration, excision and regulation of directionality (Rowley et al, 2008; McEwan, Rowley and Smith, 2009; Pokhilko et al., 2016). Research showed that mutations in the CC region allowed integrase to be active on *attL* x *attR* recombination in the absence of RDF (Rowley et al., 2008) and that the maximal efficiency of RDF is reached in a 1:1 ratio with the integrase or higher (Pokhilko et al., 2016). Moreover, it was shown that the RDF could bind the integrase in the absence of DNA (Khaleel et al., 2011). However, nothing is currently known about the site of interaction between the RDF and the coiled-coil motifs of the Int and there are yet no crystallographic structures of the Int-RDF interaction, making it a point of interest for research.

In this study, we aimed to investigate the region on the coiled-coil domain of ØC31 integrase to which its cognate RDF (gp3) might bind. Integrase mutants that were fused to gp3 were created and the ones that could not recombine *attL* and *attR* but recombined *attP* and *attB* were identified and sequenced. It was not possible to identify the site of interaction due to unforeseen errors during the construction of the mutant library. However, a mutant that was catalyzing a more unidirectional *attL* x *attR* recombination reaction with reduced *attP* x *attB* recombination was identified, and the amino acid substitutions P398L and D595N in the zinc ribbon domain were found.

## MATERIALS AND METHODS

### Strains and plasmids

*E. coli* strains used were DS941 (*recF lacI*^*q*^*lacZΔM15 argE3Δ* (*gpt-proA*) *62 his-4 leuB6 thr-1 thi-1 ara-14 lacY1 galK2 mtl-1 xyl-5 kdg K51 supE44 rpsL31 tsx-1)* (Summers and Sherratt, 1988) and DH5α (F− λ– *Δ*(*lacZYA*-*argF*)U169 *recA1 endA1 hsdR17* (r_K_ –, m_K_ +) *phoA supE44 thi*-1 *gyrA96 relA1)* (Grant et al., 1990). The plasmids used were pFEM141, pFEM33, pFM152, pFM154 and pFM211 (Figure 3), which were kindly donated by Femi J. Olorunniji of the Hooker Laboratory, University of Glasgow.

**Figure 3.**
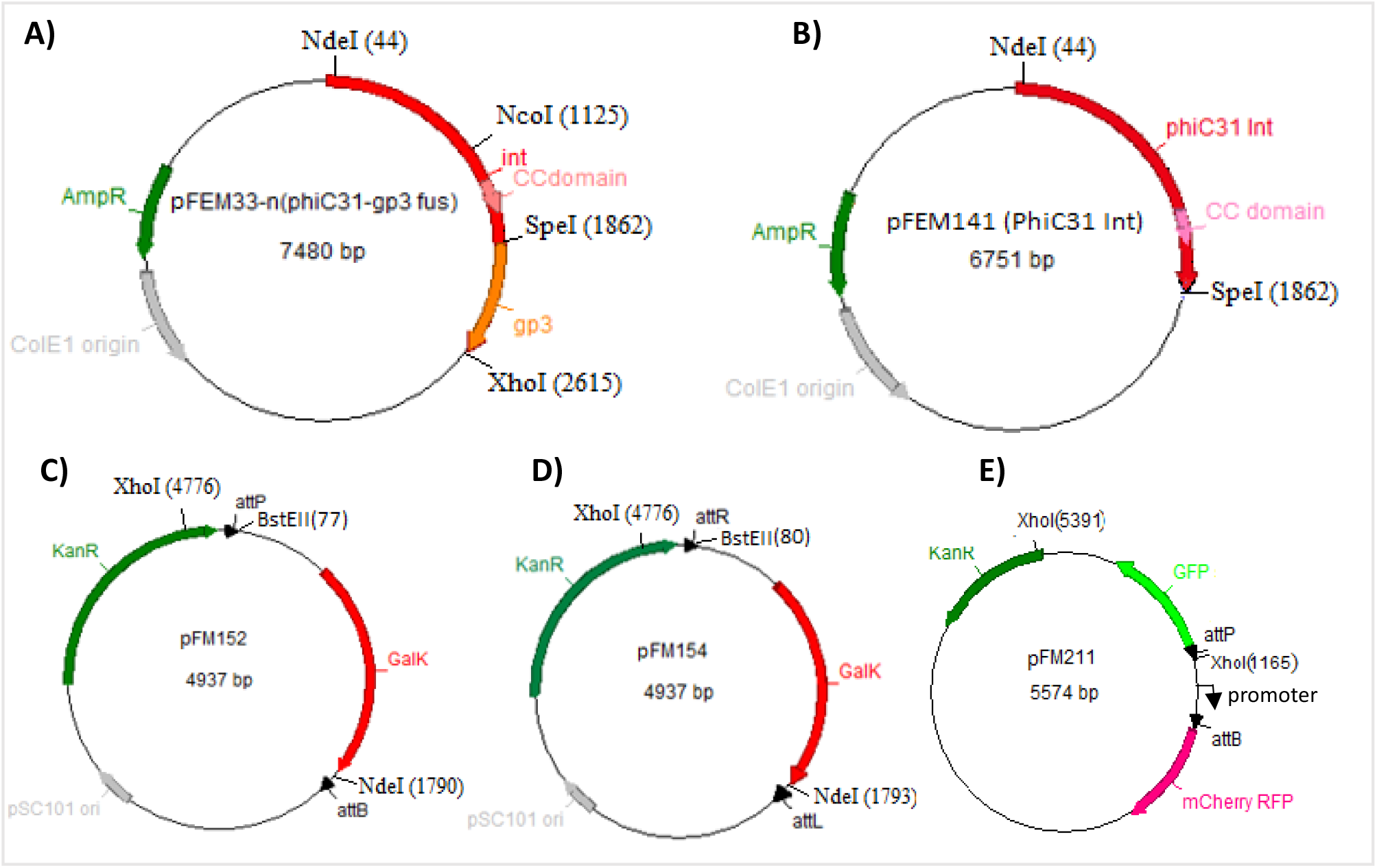
Maps of the plasmids used in this study. Ampicillin (AmpR) and kanamycin (KanR) resistance genes are indicated in green, origin of replication is shown in grey and the unique restriction sites with their position (bp) are indicated by lines. **A**. Plasmid containing the ØC31 integrase gene (red) fused to the gp3 gene (orange). **B**. Plasmid containing the ØC31 integrase gene (red). The portion of the gene encoding the coiled-coil domain (CC) of the integrase is shown in pink. **C**. Plasmid containing the galactokinase (GalK) gene (red) flanked by the attachment sites *attP* and *attB* (head-to-tail orientation) (black). **D**. Plasmid containing the GalK gene (red) flanked by the attachment sites *attR* and *attL* (head-to-tail orientation) (black). **E**. Plasmid containing the GFP (green) and RFP (pink) genes with the *attP* and *attB* sites in a head-to-head orientation (black).

### Bacterial growth conditions

Cultures were grown in Luria–Bertani (LB) broth medium or LB with 18g/l agar for plates at 37 °C overnight. The final concentrations of antibiotics used were 100 μg/ml for ampicilin and 50 μg/ml for kanamycin as required, for plasmid selection. MacConkey-galactose plates were made using MacConkey agar base (Difco) according to the manufacturer’s instructions and supplemented with 1% galactose, ampicillin and kanamycin.

### Mutagenesis and construction of mutant libraries

3 libraries of random mutants of the integrase-gp3 plasmid (pFEM33) were constructed by polymerase chain reaction (PCR) mutagenesis. DNA fragments encoding the putative coiled-coil region of the Int (region between the NcoI and SpeI of pFEM33 (see Figure 3A)) were amplified by 3 different PCR conditions using pFEM33 (Figure 3A) as template. The primers used for PCR-1 and PCR-2 were the forward primer phiC31CCmut_F; 5’AAAAAAAACGCCATGGACAAGCTGTAC 3’ that anneals to the NcoI site of pFEM33, and the reverse primer phiC31CCmut_R; 5’AAAAAAATCTACTAGTCGCCGCTACGTC 3’ that anneals to the SpeI site of pFEM33. The primers used for PCR-3 were the forward primer MHC31F; 5’ CCGGTCGAGCTTGATTGC 3’ that anneals upstream the NcoI sites, and the reverse primer MHC31R; 5’CGCTCATTCGTCTCGCTGTA 3’ that anneals downstream the SpeI site. All primers were synthesized by Integrated DNA Technologies, Inc. All PCR reactions had a final volume of 50µL with 1µM each primer and 0.08 ng/µl pFEM33 template. PCR-1 was carried out using 2X Taq polymerase mastermix (New England BioLabs) while PCR-2 and PCR-3 were carried out using dATP, dTTP, dCTP of 0.2 mM and dGTP of 0.04 mM, Taq polymerase (5U/µl) (Invitrogen) and 10X thermopol buffer (New England BioLabs). Pure water was added for a final volume of 50µL. The PCR programme was set at 95°C for denaturation (20 s), 55°C annealing (30 s), and 68°C extension (50 s for PCR-1 and PCR-2 but 60 s for PCR-3) for 30 cycles. PCR reactions were subjected to gel electrophoresis, followed by gel purification of the PCR amplicons. The gel-purified PCR amplicons and pFEM33 were subjected to restriction digest with NcoI and SpeI. The digests were subjected to gel electrophoresis followed by gel purification of the digests. Gel-purified NcoI/SpeI cut PCR amplicons were ligated to gel-purified NcoI/SpeI cut pFEM33. The ligation products were introduced into DH5α cells by CaCl_2_ transformation. The transformed cells were grown on LB plates with ampicillin at 37°C overnight. Afterwards, 900µl LB broth was added to the plates, the colonies were scrapped with a spreader and the liquid was transferred to a tube containing 20ml LB and put at 37°C overnight. Plasmid DNA from the pooled colonies was prepared and the mutant libraries were obtained. Afterwards, the extracted plasmids were screened for loss of-function or gain-of-function phenotypes as described in the Results section. Mutations were identified through sequencing, done by Eurofins Genomics.

### Transformation

DS941 and DH5α cells, with or without plasmids, were made competent and transformed following same protocol. 0.2 ml of an overnight culture was added to 10 ml of LB with ampicillin and/or kanamycin to select for plasmids (if present), and incubated at 37°C for 90 min. 1ml of culture was then transferred to a fresh Eppendorf tube for individual transformation and the cells were pelleted by centrifugation at 8000 rpm for 3.5 min at 4°C. The pellet was re-suspended in 1ml pre-chilled 50 mM CaCl_2_ followed by centrifugation at 8000 rpm for 3.5min at 4°C. Again, the cell pellet was re-suspended in 80µl CaCl_2_ and kept on ice for 1 hour. 1µl of plasmid DNA was added to the competent cells and left for 15min. The cells were heat-shocked at 37°C for 5min then put back in ice for a further 5min. 900µl LB was then added and the cells were incubated in a shaker at 37°C for 1 hour. The transformants were plated onto LB plates or MacConkey plates and incubated overnight at 37°C.

### DNA extraction and general cloning procedures

Plasmid DNA was purified from cells using a plasmid miniprep kit (Qiagen) according to the manufacturer’s instructions. Endonuclease restriction digests were done to a 20µl volume (5µl plasmid DNA, 2µl 10X cutsmart buffer (New England BioLabs), 0.5µl enzyme and pure water) for 2 hours at 37°C. Ligations were carried out at 10µl volume (4.5µl gel-purified digested PCR amplicon, 4.5µl gel-purified digested pFEM33, 1X T4 ligase buffer and 0.2U/µl T_4_ ligase), overnight at room temperature.

### Agarose gel electrophoresis and DNA gel purification

Medium-sized 1% (w/v) and 0.8% (w/v) agarose gels were run with 1X TAE buffer as per standard protocol (Sambrook and Russell, 2001). Gels were ran at 80V and 100mA for 120min. For visualization, the gels were stained with ethidium bromide (0.5 µg*/*ml), destained in demineralized H_2_O for 10 min, and imaged using Bio-Rad GelDoc UV Transilluminator. DNA purification from the gel was done using a gel extraction kit (Qiagen) according to the manufacturer’s instruction. Quantitation of bands on the gels was done as described by Pokhilko et al. (2016).

### Fluorescence assay

To measure GFP and RFP fluorescence emitted by DS941 cells containing pFM211 with or without the integrase expression plasmid, 300µl of overnight cultures were centrifuged for 3 min at 8000 rpm at room temperature, the pellets were suspended in 1.5ml phage buffer and 200µl was transferred to a 96 well flat-bottom plate. Plates were scanned using Typhoon FLA9500 scanner in fluorescence mode set to detect RFP (532nm excitation wavelength and 575nm LPG emission filter) and GFP (473nm excitation wavelength and 530DF20 BPB1 emission filter). The fluorescent intensities of RFP and GFP were quantitated using Quantity One software (Biorad) volume analysis tool. Optical density, as a measure of cell mass, was determined on a spectrophotometer (Biorad). The GFP and RFP raw fluorescence data (a.u) obtained was normalized by dividing with the transformed cells’ respective OD_600,_ and then the fluorescence output of DS941 cells was subtracted from each transformant fluorescence output.

## RESULTS

### Wild-type ØC31 Int and ØC31 Int-gp3 activity *in vivo*

To verify that *attL* x *attR* recombination by the ØC31 Int occurs only when its RDF (gp3) is present, the recombination activity of the wild-type plasmid pFEM33 encoding ØC31 Int-gp3 fusion protein (here simply referred to as Int-gp3) and that of pFEM141 encoding ØC31 Int protein (referred to as Int) was tested as described by Olorunniji et al. (2017). *E. coli* cells were first transformed with a substrate plasmid; either pFM152 that contains the *galK* gene flanked by *attP* and *attB* sites (Figure 3C) or pFM154 that contains the *galK* gene flanked by *attL* and *attR* sites (Figure 3D), and then re-transformed with either pFEM33 (Int-gp3) or pFEM141 (Int). Recombination was indirectly detected by visual inspection of the colony color present on MacConkey-galactose indicator plates (Figure 4A). Recombination of the *att* sites of either pFM152 or pFM154 results in white colonies due to the deletion of their *galK* gene (Olorunniji et al., 2017), while no recombination results in red colonies since their *galK* gene is still present. As expected, efficient recombination between *attL* and *attR* sites of pFM154 occurred in cells expressing Int-gp3 whereas no recombination occurred in cells expressing Int (Figure 4A). Moreover, recombination of *attP* and *attB* sites of pFM152 occurred in cells expressing either Int or Int-gp3 (Figure 4A).

**Figure 4.**
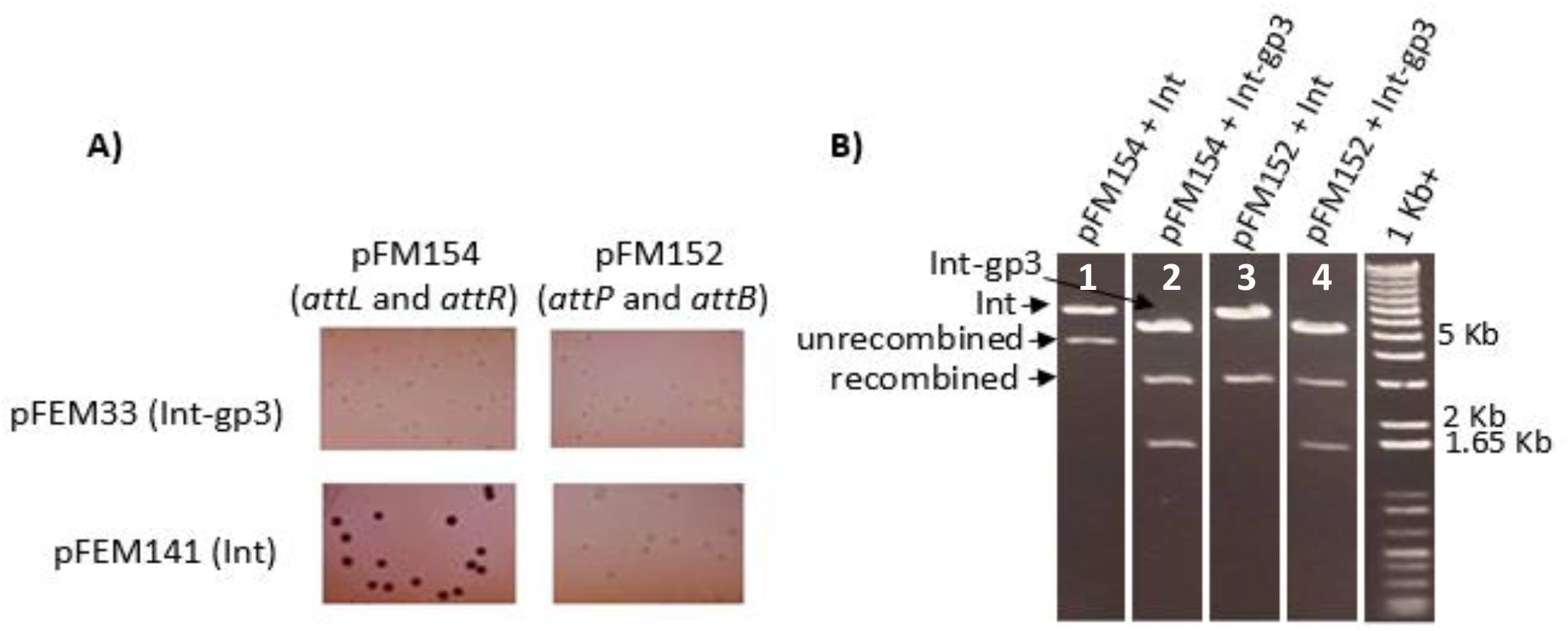
*In vivo* recombination of the substrate plasmids by the integrase and the integrase-gp3 fusion. **A)** Colony color assay; 100µl of each transformed DS941 cells was plated on MacConkey plate with 1% galactose and 150µg/ml ampicillin and 50 µg/ml kanamycin. **B)** Agarose gel electrophoresis of XhoI/BsteII digested plasmid DNA that was recovered from either a white or red colony indicated in Figure 4A. The gel (0.8% agarose) was stained with ethidium bromide and visualized under UV light.

To verify that substrate plasmids pFM152 (attP attB) and pFM154 (attL attR) were recombined in white colonies but not in red colonies, plasmid DNA was recovered from a white and a red colony, cut with XhoI and BsteII and subjected to gel electrophoresis (Figure 4B). When cut with XhoI and BsteII, pFM33 is cut twice (5.8 and 1.64 Kb fragments), pFM141 is cut only by BsteII (6.75Kb fragment), unrecombined pFM154 and pFM152 are cut twice (4.7Kb and 0.2Kb fragments) but the recombined ones are cut only by XhoI due to the deletion of the *galK* gene and loss of BsteII site (3.1 Kb fragment). The substrate plasmids pFM154 and pFM152 were recombined in cells expressing either Int or Int-gp3 (Figure 4B, lane 2 to 4) but were not recombined in cells expressing Int (Figure 4B, lane 1). Hence, *attL* x *attR* recombination by the Int only occurred when gp3 was present and the recombination activity of the wild-type integrase expression plasmids was verified.

### Integrase mutants that are insensitive to gp3

To identify the region on the coiled-coil domain of Int to which gp3 binds, mutant versions of pFEM33 (Int-gp3) that had mutations in the CC region of the Int were created. Mutant plasmids encoding defective Int-gp3 that are incapable of recombining *attL* and *attR* but still capable of recombining *attP* and *attB*, were identified using MacConkey-galactose indicator plates. This selection was done since the inability of the mutants to recombine *attL* and *attR* in the presence of RDF indicates that the Int-RDF interaction was altered and makes it possible to identify the Int-RDF site of interaction. Briefly, 3 libraries of Int-gp3 mutants were constructed through 3 different PCR conditions where the NcoI/SpeI cut PCR amplicons that encode mutated versions of the CC domain and part of the integrase (the region between the NcoI and SpeI site of pFEM33 (see Figure 3A)) were ligated to NcoI/SpeI cut pFEM33 (material and methods). Int-gp3 mutant library created using PCR-1 is referred to as mutant library 1, similarily for mutant library 2 (by PCR-2) and mutant library 3 (by PCR-3). Plasmid DNA from each mutant library was introduced into DS941 cells containing the substrate plasmid pFM154 (*attL attR*), to screen for Int-gp3 mutants with inability to recombine *attL* x *attR*. Around 80-90% of the transformants with plasmid DNA from each library, plated on MacConkey containing 1% galactose, ampicillin and kanamycin, appeared red, suggestive of defective Int-gp3 (Table 1). To verify that these mutants were unable to recombine *attL* x *attR* due to their inability to respond to gp3 and not due to complete loss of recombination function, the red colonies were taken from the plates and pooled for each mutant library separately. Plasmid DNA was prepared from the pooled colonies and introduced into DS941 cells containing pFM152 (*attP attB*) to screen for the mutants that were still capable of recombining *attP x attB*. Around 20-30% of the transformants appeared white, indicative of Int-gp3 mutants that can recombine *attP* x *attB* while rendered insensitive to gp3 and unable to recombine *attL* x *attR*. Therefore, 10 Int-gp3 mutants were randomly selected from each library (30 mutants in total) and re-transformation assays were done as described next.

**Table 1.** Percentage of DS941 cells that appeared red with pFM154 (*attL attR*) and white with pFM152 (*attP attB*) when expressing mutant Int-gp3 on MacConkey plates.

| Mutant library | Number of mutants | % Red with *attL* and *attR* | % White with *attP* and *attB* |
| --- | --- | --- | --- |
| 1 | 154 | 94% | 28% |
| 2 | 168 | 79% | 19% |
| 3 | 113 | 92% | 17% |

### *In vivo* recombination with defective Int-gp3

To test the 30 selected Int-gp3 mutants for inability to recombine *attL* and *attR* but efficiently recombine *attP* and *attB*, re-transformation assays with DS941 cells containing either the plasmid substrates pFM152 (*attP attB*) or pFM154 (*attL attR*) were done and recombination was indirectly detected as mentioned before. Out of the 30 selected mutants, 3 mutants from mutant library 1 (designated Int-gp3^1^, Int-gp3^2^ and Int-gp3^3^) and 1 mutant from mutant library 2 (designated Int-gp3^4^), showed low levels of *attL* x *attR* recombination of pFM154 (mostly red colonies) and a high level of *attP x attB* recombination of pFM152 (white colonies), compared to the wild-type Int-gp3 that recombined both pFM152 and PFM154 (Figure 5).

**Figure 5.**
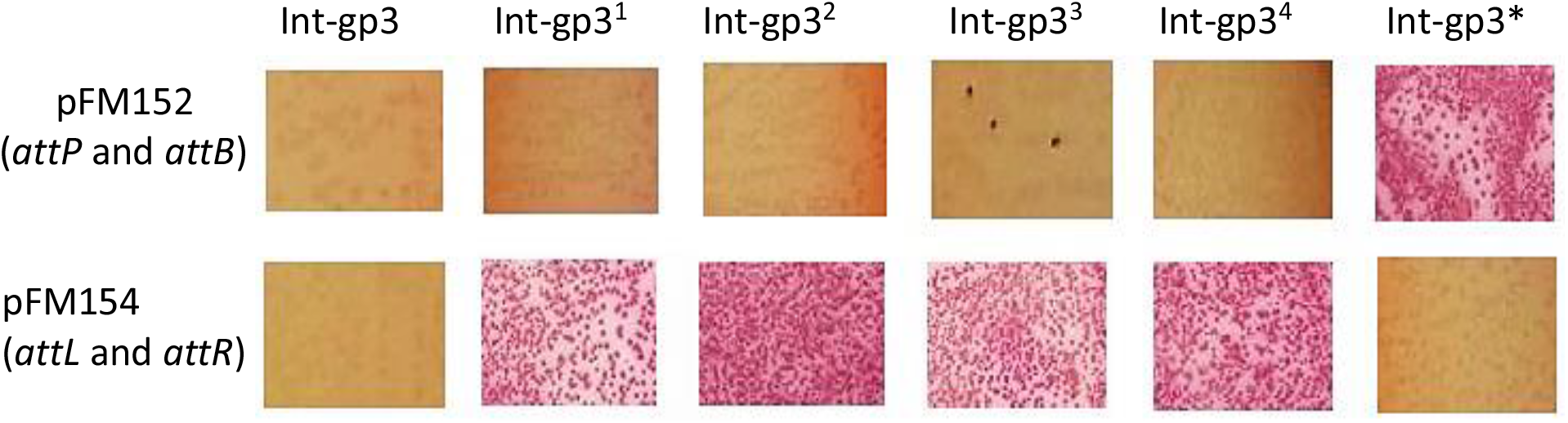
*In vivo* recombination with the gp3-insensitive integrase. DS941 cells with either pFM152 or pFM154 were transformed with wild-type and defective Int-gp3. 100µl of each transformed DS941 cells were plated on MacConkey plate with 1% galactose and 150µg/ml ampicillin and 50 µg/ml kanamycin. Recombinants were detected indirectly by deletion of *galK* (white colonies) flanked by two *att* sites in pFM152 and pFM154.

Moreover, a single mutant (designated Int-gp3*) from mutant library 3, showed a high level of *attL* x *attR* recombination of pFM154 (white colonies) with reduced level of *attP x attB* recombination of pFM152 (red colonies) compared to the wild-type Int-gp3 (Figure 5). Int-gp3* was selected since it suggested that it was catalyzing a more unidirectional attL x attR recombination reaction compared to the wild type Int-gp3 (ref). The remaining mutants had less efficiently recombined *attL* x *attR* and *attP* x *attB* of pFM154 and pFM152 respectively and the resulting colonies were a mix of white and red.

To analyze the recombination products, plasmid DNA was prepared from the colonies on the plate and analyzed by agarose gel electrophoresis (Figure 6). Wild-type Int-gp3 mediated recombination of *attL* x *attR* test substrate pFM154 and *attP* x *attB* test substrate pFM152 results in the deletion of the *galK* gene that alters the size of the plasmid, resulting in a distinct band that runs faster on the gel (Figure 6, lanes 2 and 9) compared to that of an unrecombined plasmid (Figure 6, lane 1 and 8).

**Figure 6.**
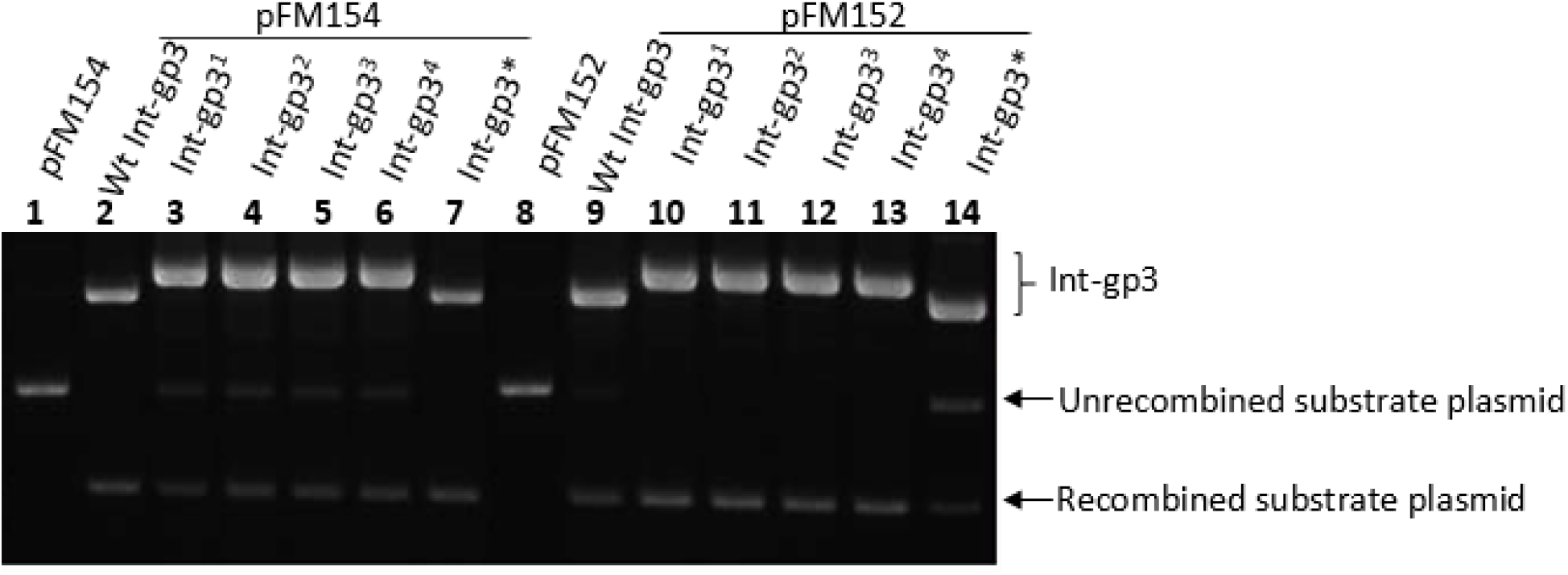
Agarose gel electrophoresis of uncut wild-type and mutant Int-gp3 with the substrate plasmids. The gel (0.8% agarose) was stained with ethidium bromide and visualized under UV light. The wild-type (Wt) and mutant Int-gp3 plasmids were present with either pFM152 or pFM154 as indicated.

In agreement with the observed colony colors (Figure 5), the *attL* x *attR* test substrate pFM154 was mostly unrecombined in cells expressing Int-gp3^1^, Int-gp3^2^, Int-gp3^3^ and Int-gp3^4^ (Figure 6, lanes 3 to 6), whereas it was completley recombined in cells expressing Int-gp3* (Figure 6, lane 7).

However, the *attP* x *attB* test substrate pFM152 was completely recombined in cells expressing Int-gp3^1^, Int-gp3^2^, Int-gp3^3^ and Int-gp3^4^ (Figure 6, lane 10 to 13), but was mostly unrecombined in cells expressing the Int-gp3*(Figure 6, lane 14). Unexpectedly, Int-gp3^1^, Int-gp3^2^, Int-gp3^3^ and Int-gp3^4^ expression plasmids were bigger in size than the wild-type Int-gp3 (Figure 6, compare lanes 3-6 with lane 1and lanes 10-13 with lane 8).

### Sequencing of the defective Int-gp3 mutants

The mutant plasmids, Int-gp3^1^, Int-gp3^2^, Int-gp3^3^, Int-gp3^4^ and Int-gp3*, were sequenced using the PCR primers MHC31F (upstream NcoI site) and MHC31R (downstream SpeI site), in order to identify the mutations responsible for the previously observed phenotypes (see Figure 5). The mutants, Int-gp3^1^, Int-gp3^2^, Int-gp3^3^ and Int-gp3^4^ contained two copies of the PCR amplicon inserted between the NcoI and SpeI sites of pFEM33 (Figure 7), consistent with the large size seen on the gel (Figure 6). This duplication of the 737bp PCR product caused a frame-shift mutation in the C-terminal domain of the integrase, resulting in a stop codon downstream the first NcoI/SpeI PCR fragment (Figure 7). Translation results in a full Integrase but no gp3. Therefore, these mutants showed no *attL* x *attR* recombination due to the lack of gp3 proteins that resulted in the observed red colonies, but did not affect their attP x attB recombination ability, resulting in white colonies (Figure 5). Hence, it was not possible to identify the region on the coiled-coil domain of ØC31 Int to which gp3 binds.

**Figure 7.**
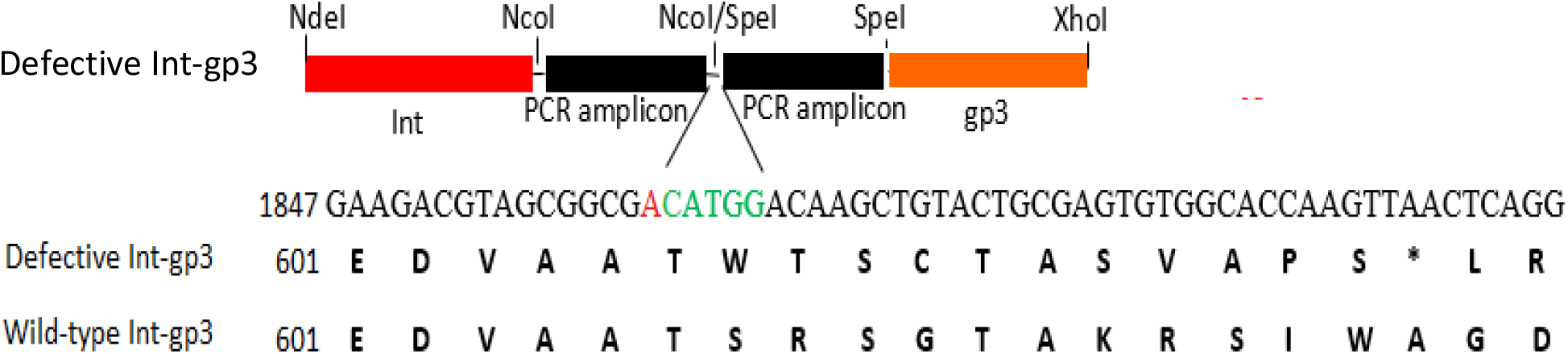
Schematic representation of the Int-gp3 portion of the mutant Int-gp3 expression plasmid. The gene encoding the integrase is shown in red, the two copies of PCR amplicon in black and the gp3 in orange. The restriction sites are as indicated. Part of the DNA sequence (1847-1907 bp) from the sequenced Int-gp3 mutant is shown with the NcoI/SpeI junction colored red (SpeI) and green (NcoI). The amino acid sequence translated from the top DNA sequence of the mutant Int-gp3 is shown, as well that of the wild-type Int-gp3. * represent a stop codon.

However, Int-gp3* had 2 point mutations that caused P398L substitution in the zinc ribbon domain of the Int and D595N substitution near its end. These mutations in the Int-gp3* affected its structure, making it less capable for *attP* x *attB* recombination which resulted in the observed red colonies, but did not affect its *attL* x *attR* recombination activity which resulted in the observed white colonies (Figure 5).

### Characterization of Int-gp3*

To test if the Int-gp3* was catalyzing a more unidirectional *attL* x *attR* recombination reaction with a much lower level of *attP* x *attB* recombination than the wild-type Int-gp3, its recombination activity was tested with an inversion plasmid pFM211 that contains head-to-head orientation of *attP* and *attB* sites and encodes the fluorescence genes GFP and RFP (Figure 3E). When no recombination occurs, the inversion plasmid pFM211 generates RFP fluorescence whereas recombination of its *att* sites flips the orientation of transcription (Olorunniji et al., 2017) resulting in GFP fluorescence (see Figure 3E).Therefore, GFP fluorescence acts as a reporter of *attP* x *attB* recombination. DS941 cells with PFM211 were transformed with Int-gp3* and Int-gp3 expression plasmids and the RFP/GFP fluorescence of 3 replicates and the controls were measured using Typhoon FLA9500 scanner (Figure 8).

**Figure 8.**
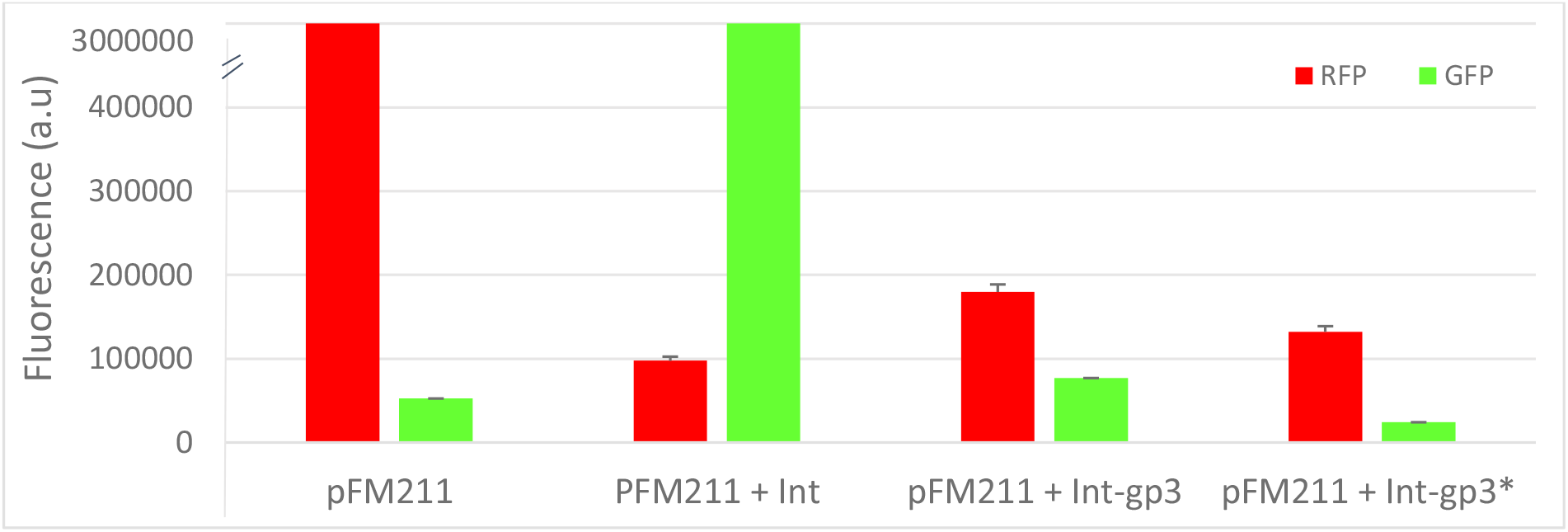
Fluorescence of the expressed RFP and GFP proteins of pFM211 in the presence/absence of Int and Int-gp3 in DS941 cells. Fluorescence of 200μl culture each present in a 96-well plate was measured using Typhoon FLA9500 scanner. The raw data was handled as present in the method section and this plot was made. The data are the average and standard error for three replicates.

As expected, efficient recombination of pFM211 occurred in cells expressing the Int (GFP fluorescence) whereas a much lower level of recombination was observed in cells expressing the wild-type and mutant Int-gp3 (Figure 8). Moreover, DS941 cells expressing Int-gp3* generated a lower GFP fluorescence output compared to the cells expressing Int-gp3 (Figure 8), indicating that the mutant was less active on *attP* x *attB* recombination and indirectly suggesting the possibility of a more unidirectional attL x attR reaction, but the Int-gp3* was not tested on an inversion plasmid with *attL* and *attR* sites.

RFP fluorescence was observed in all cells, but with higher output by cells expressing the mutant and wild-type Int-gp3 compared to the ones expressing the Int (Figure 8), indicating a higher level of unrecombined pFM211.

In order to quantify the relative amounts of the unrecombined and recombined pFM211 in these cells, plasmid DNA was isolated, subjected to restriction digest with XhoI (but for the Int expression plasmid+pFM211, EcoRI and XhoI were used) followed by gel electrophoresis and the obtained bands were quantified (Figure 9).

**Figure 9.**
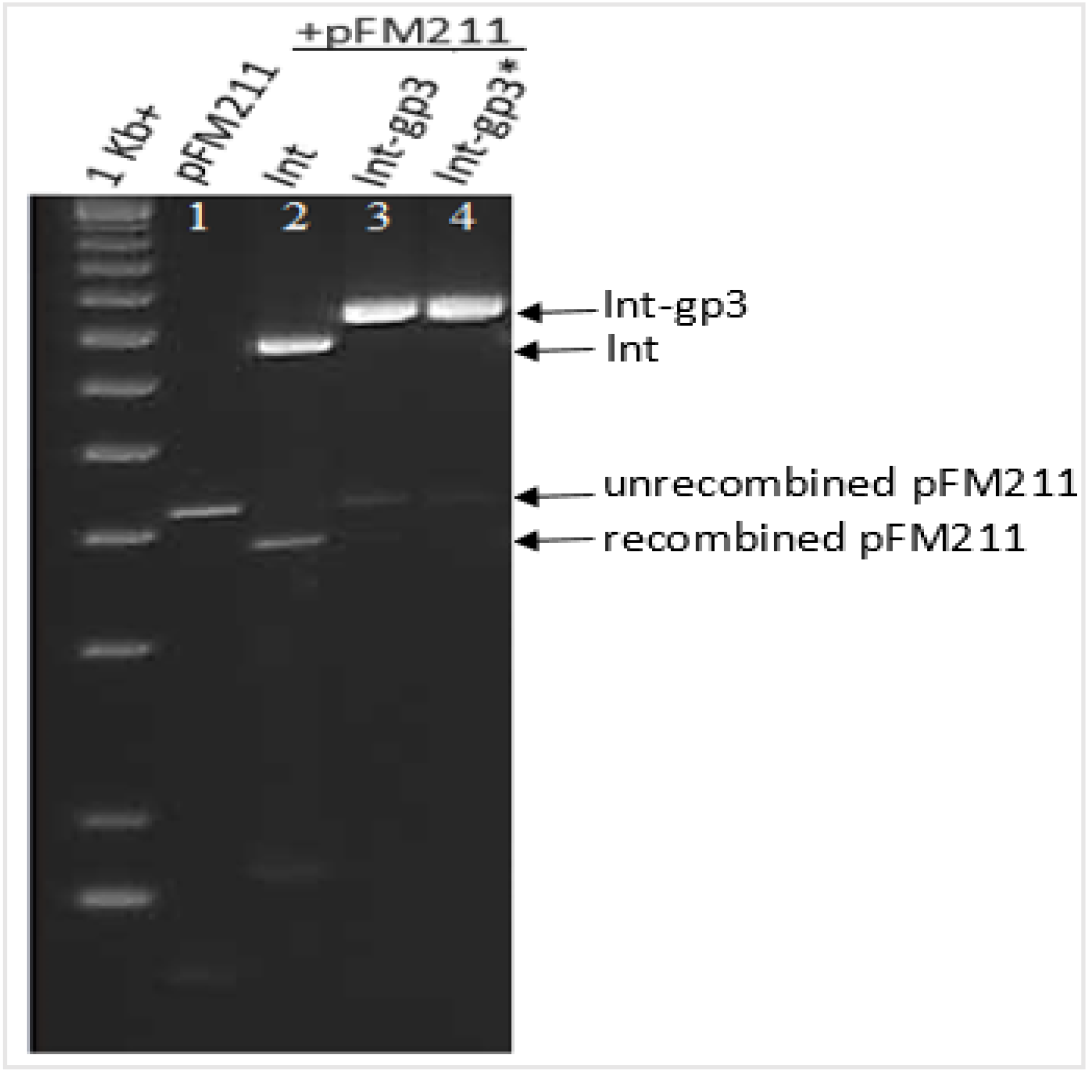
Agarose gel electrophoresis of digested plasmid DNA recovered from DS941 cells with pFM211 and expressing either wild-type or mutant Int-gp3 or Int. The gel (1.2 % agarose) was stained with ethidium bromide and visualized under UV light. All plasmids were digested with XhoI except for the Int+pFM211 that was digested with XhoI and EcoRI.

Recombination of pFM211 does not alter the size of the plasmid but changes the sizes of the restriction digest products (Olorunniji et al., 2017), resulting in distinct DNA fragments from unrecombined and recombined pFM211 molecules when cut with XhoI. When cut with XhoI, pFEM33 is cut once (7.48Kb fragment), unrecombined pFM211 is cut twice (4.2 Kb and 1.34Kb fragments), recombined pFM211 is cut twice (3.7 Kb and 1.8Kb fragments) and pFEM141 is only by EcoRI once (6.75 Kb fragment). As expected, the inversion plasmid pFM211 (attP attB) was recombined efficiently (100%) in cells expressing Int (Figure 9, lane 2), but it was poorly recombined (52%) in cells expressing Int-gp3 (Figure 9, lane 3). Moreover, pFM211 was less efficiently (28%) recombined in cells expressing Int-gp3*. Therefore, Int-gp3* was catalyzing a more unidirectional *attL* x *attR* recombination reaction with a much lower level of *attP* x *attB* recombination than the wild-type Int-gp3.

## DISCUSSION

We described here experimental approaches that aimed to identify the possible region to which the RDF might bind on the integrase, by creating integrase mutants (with possible mutations in their coiled-coil domain) fused to gp3 (RDF), and selecting for mutants that were inactive for *attL* x *attR* recombination but not *attP* x *attB* recombination. Similar approaches have been used in understanding the control of directionality in integrases (Rowley et al., 2008; Liu et al., 2010). Through sequencing, the mutations that caused the integrase inactivity for *attL* x *attR* recombination in the presence of RDF could be identified and the integrase-RDF binding region could be recognized. However, we were not able to characterize the Int-gp3 mutants and identify the binding region.

The selected Int-gp3 mutants, Int-gp3^1^, Int-gp3^2^, Int-gp3^3^ and Int-gp3^4^ that showed to be inactive on *attL* x *attR* recombination but were active on *attP* x *attB* recombination (Figure 5) had given false positive results. The mutant Int-gp3 expression plasmids encoding these Int-gp3 mutants were constructed by ligating NcoI/SpeI cut PCR amplicons (encoding mutated versions of the CC domain) to NcoI/SpeI cut wild-type Int-gp3 plasmid. Unexpectedly, the mutant plasmids were bigger in size compared to the wild-type Int-gp3 expression plasmid. This size difference was due to the presence of two copies of PCR amplicons in the mutant plasmids which ligated to each other at the NcoI/SpeI restriction site of one another. This occurred due to the unexpected compatibility of the NcoI and SpeI restriction sites, where the nucleotides C and G within the NcoI site in one amplicon had bind to the G and C nucleotides within the SpeI sites of another amplicon (see Figure 10). The resulting ligation product shown in Figure 10 was consistent with the sequencing results of the Int-gp3 mutant plasmids (Figure 7). To avoid this from happening, different restriction enzymes could be used.

**Figure 10.**
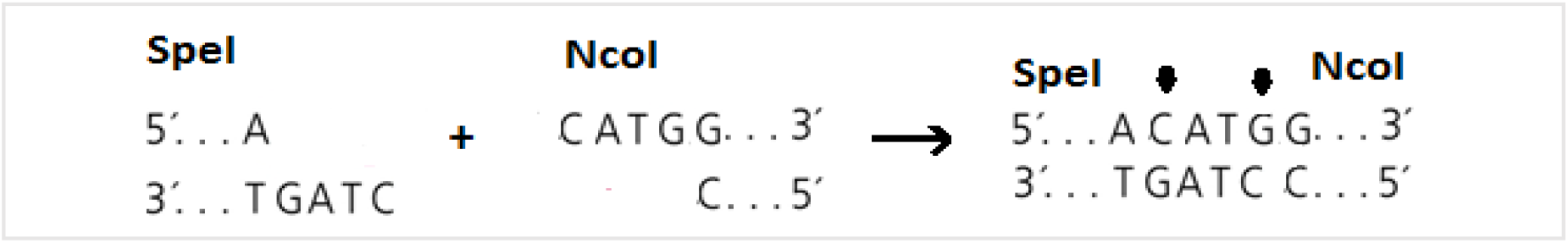
Illustration showing the ligation between NcoI and SpeI sites. The black arrows indicate the complementary binding of the nucleotides responsible for the unexpected compatibility between NcoI and SpeI site.

Moreover, the remaining mutants that were not efficiently recombining *attL* x *attR* and *attP* x *attB* and resulted in a mix of white and red colonies were due to an experimental error while selecting for the mutants. Two colonies were picked by mistake and the resulting plasmid DNA was a mix of wild-type and defective Int-gp3.

The mutations (P398L and D595N) in the Int-gp3* that conferred its ability for a more unidirectional *attL* x *attR* recombination with reduced *attP* x *attB* recombination lay within the zinc ribbon domain of the mutant integrase. Previous work has shown that the zinc ribbon domain of the integrase is required for high affinity binding of the integrase to the *attP* and *attB* sites (Ghosh, Pannunzio and Hatfull, 2005) and mediates the interactions between the integrase subunits (Van Duyne and Rutherford, 2013). As well, it is proposed that the recombination efficiency is mainly affected by the decrease of binding affinity (Liu et al., 2010). Therefore, the mutations that occurred in the zinc ribbon domain of the Int-gp3* had affected the integrase interaction with the *attP* and *attB* sites, resulting in the reduced level of *attP* x *attB* recombination. Similar results were observed in a study by McEwan et al. (2011) were mutations in this region affected integrase interaction with both *attP* and *attB* sites. Moreover, a similar integrase mutant with the amino acid substitution Y393A in the zinc domain had completely lost the ability to bind *attB* (Liu et al., 2010). Furthermore, these mutations in the Int-gp3* enhanced its ability to recombine *attL* and *attR* resulting in a more unidirectional reaction. The reaction catalyzed by Int-gp3* had similar features to Bxb1 integrase which catalyzes *attL* x *attR* recombination in the presence of its cognate RDF, gp47 (Ghosh, Bibb and Hatfull, 2008). Moreover, when the Int-gp3* was tested for unidirectionality of *attL* x *attR* recombination, its activity was only tested on an inversion plasmid with *attP* and *attB sites*. However, for better accuracy in testing its unidirectionality, the activity should have been tested on an inversion plasmid with *attL* and *attR* sites. It was not possible to do so due to the unavailability of this plasmid and time constraints of the project. If more time was allowed, the inversion plasmid with *attP* and *attB* could be recombined by integrase to generate an inversion plasmid with *attL* and *attR* sites which could be isolated and tested with the Int-gp3*. Furthermore, RFP fluorescence from the inversion plasmid pFM211 was extremely high when unrecombined because of the aggregation of RFPs (Piatkevich and Verkhusha, 2011). This mutant could be used in synthetic switches for the creation of biological computers that count and record stimuli (Fogg et al., 2014; Baumgardner et al., 2009).

To further improve this work a new library of Int-gp3 mutants could be made and screened for by following the same experimental approaches described here, but using different restriction enzymes when constructing the mutant plasmids. This would allow the identification of the Int-RDF binding region. In addition, the wild-type Int-gp3 fusion protein could be purified and the structure of the Int-RDF interaction could be identified by employing X-ray crystallography since no structures are yet available. This paper now provides approaches towards understanding the Integrase-RDF interaction.

## Abbreviations used

Int: Integrase
Int-gp3: Integrase-gp3
RDF: Recombination directionality factor
CC: Coiled-coil
GFP: Green Fluorescent Protein
RFP: Red Fluorescent Protein

## Notes

### Competing Interest Statement

The authors have declared no competing interest.

### Summary of Updates

The text in the abstract was updated to include a reference to this paper: Paul C M Fogg, Ellen Younger, Booshini D Fernando, Thanafez Khaleel, W Marshall Stark, Margaret C M Smith, Recombination directionality factor gp3 binds ϕC31 integrase via the zinc domain, potentially affecting the trajectory of the coiled-coil motif, Nucleic Acids Research, Volume 46, Issue 3, 16 February 2018, Pages 1308-1320, https://doi.org/10.1093/nar/gkx1233

